# Anterior cingulate cortex connectivity is associated with suppression of behavior in a rat model of chronic pain

**DOI:** 10.1101/225482

**Authors:** Laurel S. Morris, Christian Sprenger, Ken Koda, Daniela M. de la Mora, Tomomi Yamada, Hiroaki Mano, Yuto Kashiwagi, Yoshichika Yoshioka, Yasuhide Morioka, Ben Seymour

**Author notes:** LM, CS, KK contributed equally to this study.

## Abstract

A cardinal feature of persistent pain that follows injury is a general suppression of behavior, in which motivation is inhibited in a way that promotes energy conservation and recuperation. Across species, the anterior cingulate cortex (ACC) is associated with the motivational aspects of phasic pain, but whether it mediates motivational functions in persistent pain is less clear. Using burrowing behavior as an marker of non-specific motivated behavior in rodents, we studied the suppression of burrowing following painful CFA or control injection into the right knee-joint of 37 rats (18 with pain), and examined associated neural connectivity with ultra-high-field resting state functional MRI. We found that connectivity between ACC and the subcortex correlated with the reduction in burrowing behavior observed following the pain manipulation. In a full replication study we confirmed these findings in a group of 44 rats (23 with pain). Across both datasets, reduced burrowing was assosciated with increased connectivity between ACC and subcortical structures including hypothalamic/preoptic nuclei and the bed nucleus of the stria terminalis. Together, the findings implicate ACC connectivity as a robust correlate of the motivational aspect of persistent pain in rodents.

## 1 Introduction

One of the greatest challenges in the development of novel analgesics is the difficulty in evaluating pain in animal models of chronic pain. Pain is a uniquely private and subjective experience, and the mainstay of outcome measures in clinical trials remains subjective ratings. Since these are not available in animals, evaluation of pain depends on surrogate measures of behavior, primarily motor responses directly related to evoked or spontaneous pain in the region of injury. However, the difficulty in translating these behaviors to humans is well recognised, and this greatly hampers the ability to predict whether analgesics that are successful in animals will work in humans. This has led to a requirement for measures of behavior that better reflect persistent pain in animals, and which have translational validity to humans.

The advent of functional neuroimaging offers a novel approach to the evaluation of pain in animals. In humans, a broad set of cortical and subcortical regions are implicated in pain processing and the expression of pain behavior. This diversity reflects the fact that pain is a multidimensional experience, engaging sensory, affective and cognitive processing. Human studies of phasic experimental pain implicate a broad network of pain regions (the ‘pain matrix’) that includes thalamus, primary and secondary somatosensory cortex (S1, SII), cerebellum, the insula cortex, and the anterior cingulate cortex (ACC) (Moisset and Bouhassira, 2007; Price, 2000). Although the precise functions of each region remains unclear, there is a consensus that some regions (thalamus, primary and secondary somatosensory cortex, cerebellar) better reflect sensori-motor processes, and others (the anterior insula cortex and in particular the ACC) better reflect affective processes (Büchel et al., 2002; Lloyd, Di Pellegrino, and Roberts, 2004; Rainville, 2002; Rainville et al., 1997; Singer et al., 2004). However, there remains uncertainty as to how well studies of phasic experimental pain translate to the persistent pain seen in chronic pain models. Although phasic pain occurs as a part of many chronic pain conditions, as seen with spontaneous phasic and hyper-sensitive evoked pain, the nature of persistent underlying pain is quite different, and serves a distinct behavioral function to reduce activity and conserve energy.

The ACC, consisting of Brodmann area’s 24 and 25, is a strong candidate to play a dominant role in mediating the affective behavioral manifestations of persistent pain. It serves a primary function as a modulator of the affective tone of visceral, motor and endocrine efferents to downstream regions involved in responses to nociceptive stimuli, including the peri-acqueductal grey (PAG) and amygdala (Calejesan, Kim, and Zhuo, 2000a; Vogt, Finch, and Olson, 1992; Zhuo, 2007).

In persistent pain, some evidence suggests it may operate in a network with other areas, especially somatosensory (S1) cortex, in sensorimotor manifestations of pain (such as mechanical allodynia) (Eto et al., 2011), and long term hyperalgesia in response to pain conditions in rats is associated with reduced S1, ACC and insula volumes (Seminowicz et al., 2009).

Recently, novel behavioral tasks such as burrowing behavior have been proposed as a measure of the affective impact of pain. Burrowing behavior is an innate, spontaneous behavior that demonstrates the overall ‘wellbeing’ or affective-motivational tone of the rat (Deacon, 2006) and is not affected by limb hypersensitivity (Andrews et al., 2012). Burrowing provides shelter and protection from environmental predators, food storage and foraging, but also comes with energy costs (Reichman and Smith, 1990). This behavior is reduced in rats following peripheral nerve injury or pain (Andrews et al., 2012), and therefore may provide a useful index of the motivational component of persistent background chronic pain.

Taken together, these considerations suggest that changes in ACC connectivity may be related to the modulation of general motivational behaviors, such as burrowing, by persistent pain. To test this hypothesis, therefore, we studied burrowing behavior in adult rats in a model of inflammatory arthritis pain (intra-articular injection of complete Freund’s adjuvant to the hind leg), and measured ACC functional connectivity in the brain using fMRI. We performed two studies: an initial study in a 11.7T MRI scanner, and a replication study using a 7T MRI scanner, to confirm the results.

## 2 Materials and Methods

### 2.1 Experimental Animals

In experiment 1, 72 male LEW/CrlCrlj rats (Charles River Laboratories, Japan, Inc.) were used. In experiment 2 (replication study) we studied a further 144 male rats. Rats were housed in groups of 3 in plastic cages under controlled temperature and humidity, and provided free access to food and water under a 12/12 hour reversed light-dark cycle. All procedures were approved by internal animal care and use committee of Shionogi Pharmaceutical Research Center (Osaka, Japan) instructed by Association for Assessment and Accreditation of Laboratory Animal Care International (AAALAC) guidelines.

### 2.2 Preparation of pain model rats

Rats were anaesthetized with isoflurane (Mylan, Canonsburg, PA, USA) and then intra-articular injected with 25 μl of either Freund’s Complete Adjuvant (CFA; Mycobacterium butyricum [BD DIFCO, Franklin Lakes, NJ, U.S.]; 2 mg/ml of liquid paraffin [Maruishi Pharmaceutical Co., Ltd., Osaka, Japan]), or vehicle (sham rats) into the right knee joint of the hind leg. In both cases, the left hind knee joint was kept untreated. Both the pain model and the sham rats were 6-weeks-old at the time of injection.

### 2.3 Pain behavior

In experiment 1, rats were tested for pain behavior 18 and 22 days after FCA/vehicle injection. First, the knee diameter (KD) was determined using a digita caliper (CD-15CX, Mitutoyo Corporation, Kanagawa, Japan) for 22 days after FCA/vehicle injection. Second, a weight bearing difference test (WBD) on each hind leg was performed (Bioseb, Boulogne, France). Values of WBD are obtained using the following formula, Where *W_R_* corresponds to the amount of weight put on the right leg and W_L_ to the amount of weight put on the left leg. This formula expresses the percentage of the rat’s body weight put on the CFA/saline-injected right leg, thus a 50% value means equal weight distribution across both hind legs.

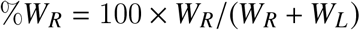

Third, a grip strength test (GS, San Diego Instruments, San Diego, CA, USA) was conducted to quantify the mechanical strength of the hind legs and hence control for the development of pain in model rats. Briefly, each rat was gently restrained and allowed to grasp the wire mesh frame with its hind limbs and was moved in a rostral-to-caudal direction until the grip released. Finally, burrowing behavior (BB) was investigated. For burrowing experiments plastic tubes (32 cm in length and 10 cm in diameter) were filled with 3 kg of gravel (5-8 mm) and placed in Plexiglas cages (560 × 440 × 200 mm). The open-end of the tube was elevated 6 cm from the floor of the cage. Rats were allowed to individually burrow during 30 min for 18 days after FCA or vehicle injections were performed and the amount of gravel burrowed was recorded. In experiment 2, rats were tested for pain behavior 20 and 21 days after FCA/vehicle injection. The KD was determined for 21 days after FCA/vehicle injection. WBD and GS was performed for 20 days after FCA/vehicle injection. In BB, baseline burrowing performance was tested on day 0 after the 2 days social facilitation. For social facilitation, rats were placed into Plexiglas cages in pairs and spent 30 min in an empty cage, after which the gravel-filled plastic tubes was introduced for a 60-min session. After the baseline burrowing test, animals were assigned to groups based on their burrowing performance and intra-articular injected FCA/vehicle. For 20 days after FCA/vehicle injection, rats were allowed to individually burrow for 30 min and the amount of gravel burrowed was recorded.

### 2.4 Animal selection and transfer to MRI facilities

In experiment 1, based on the results obtained from the WBD on the 18th day, rats were selected out of the ones that displayed either moderate or severe pain and were sent to the MRI facility. fMRI scanning was conducted at the Center for Information and Neural Networks (CiNet, Osaka University, Suita, Japan). After being transferred to the MRI facility, animals were kept for four days under standard laboratory conditions (room temperature of 22-23°C and a 12-h light/dark cycle) with free access to food and water. MRI data was acquired on the third day (i.e. 21 days after FCA/saline injection). On the fourth day, behavioral data was obtained once more. While in experiment 2, based on the results obtained from the WBD on the 20th day, rats were selected. fMRI measurements were performed at an on-site (Shionogi & Co. Ltd., Osaka, Japan) MRI facility on 21st day after FCA/vehicle injection.

### 2.5 Resting state functional MRI data acquisition

In Experiment 1 data was acquired from a total of 37 rats (18 CFA pain model) with an 11.7 Tesla Avance II vertical bore system (Bruker BioSpin, Ettlingen, Germany) and a home-made transmit/receive surface radio frequency (RF) coil. Rats were anesthetized with a mixture of air and 2.8% isoflurane (Wako Pure Chemical Industries Ltd., Osaka, Japan) and then placed in an MRI-compatible animal cradle. The isoflurane concentration was mantained at 2% ± 0.5%, adjusted to maintain the respiration rate at 70 ± 10 breaths/min throughout the sessions. An axial T2-weighted (T2W) imaging was performed using a rapid acquisition of relaxation enhancement (RARE) sequence (repetition time/echo time [TR/TE] = 6500/45 ms, number of averages [NA] = 8, field of view [FOV] = 32 × 16 mm, matrix size = 256 × 256, slice thickness = 500 μm and acquisition time = 14 min). To acquire resting state funcional MRI (rsfMRI) data we performed gradient-echo echo-planar imaging (TR/TE = 2500/7ms, number of segments = 2, Flip Angle = 60°, FOV = 51.2 × 51.2 mm, matrix size = 64 × 64, slice thickness = 1 mm, in-plane resolution = 800 × 800 *μm*^2^, bandwidth = 300 kHz, and acquisition time = 20 min).

In Experiment 2 MRI data from 44 rats (21 controls, 23 with pain) were acquired on a 7 Tesla Agilent (Varian) System (Agilent, Palo Alto, CA, USA) equipped with a two channel surface receiver coil. Directly before begin of the experiments animals were anesthetized with an initial concentration of 5% isoflurane (Pfizer Japan Inc., Tokyo, Japan) in 100% oxygen which was reduced after approximately 1 min to 2%. To minimize imaging artifacts related to magnetic susceptibility differences the middle ears were then filled with conventional ultrasound gel (Aquasonic 100, Parker Laboratories, Fairfield, NJ). During fMRI the isoflurane concentration was maintained at 1.5% ± 0. 5%. Axial T2-weighted anatomical images were acquired using likewise a fast spin-echo sequence (TR = 4000ms, ETL = 2, effective TE = 50 ms, NA = 3, FOV = 22.4 × 22.4 mm, matrix size = 128 × 128, slice thickness = 0.5 mm, acquisition time = 13 min). Resting state fMRI data were acquired employing an echo-planar gradient-echo imaging sequence collecting phase maps between repetitions (Varian’s EPIP: TR/TE 1600/18ms, Flip Angle = 90°, FOV = 22.4 × 22.4 mm, matrix size = 64x64, slice thickness = 1 mm, bandwidth = 3.9 kHz, acquisition time = 13.3 min).

### 2.6 Imaging data preprocessing

The imaging data preprocessing in experiment 1 was performed with AFNI (https://afni.nimh.nih.gov) and FSL (FMRIB, University of Oxford, UK; www.fmrib.ox.ac.uk/fsl/melodic2/index.html). Data from 7 animals could not be analyzed due to technical difficulties and resulting poor image quality leaving 30 animals for the final rsfMRI analysis. Functional data was corrected for slice timing offsets, linear detrended and corrected for movement (6 df). Functional data were up-sampled to the resolution of the anatomical images, structural images underwent skull stripping, and both were coregistered (in a similar manner to (Kundu et al., 2014)). A study-specific anatomical template was subsequently created by averaging four normal control rat anatomical datasets. Functional and anatomical data for each rat were then normalized to this study-specific template using the normalization procedure implemented in AFNI. Functional data were furthermore subjected to despiking, smoothing (full width half maximum = 0.5mm), band pass filtering and linear regression of motion parameters.

In experiment 2 imaging data preprocessing was carried out using SPM (SPM12, Wellcome Trust Centre for Neuroimaging, London, UK). Imaging data from 1 CFA treated animal could not be analyzed due to an insufficient result of the spatial normalization procedure (see below) leaving 43 animals for the final analysis. Functional data were corrected for differences in slice timing and movements (6 df). To spatially normalize the data, we first coregistered the functional data with the individual anatomical images using normalized cross-correlation as the objective function. Next, we segmented the T2-weighted anatomical images using the tissue priors published by (Valdés-Hernández et al., 2011). Subsequently, tissue probability maps were DARTEL-imported (Ashburner, 2007) and a study-specific DARTEL group template was generated. The functional data were then normalized to this study-specific DARTEL template and up-sampled to a resolution of 0.125x0.125x0.125mm3. Finally, functional data were spatially smoothed, temporally filtered and subjected to regression of movement parameters similar to experiment 1.

### 2.7 Assessment of functional connectivity

Functional connectivity (Friston, 1994) of the ACC was assessed with a seed based functional connectivity approach. The ACC seed region was determined as the anterior portion of Brodmann area 24 (Vogt and Peters, 1981), approximately 1mm rostral to Bregma (Calejesan, Kim, and Zhuo, 2000b). The mean time series in this region were extracted for each animal from a sphere of 0.5mm radius around this coordinate and functional connectivity was determined by calculating brain-wide z-transformed correlation maps based on the preprocessed functional time series. Subsequently, functional connectivity maps were compared between groups employing a two-sample t-test. Finally, correlations between the individual functional connectivity maps and the behavioral measures (KD, DWB, GS, BB) were calculated across animals and compared between groups employing likewise t-statistics. To allow for comparability of the results from experiment 1 and 2, t-maps representing the statistical results and the study-specifc anatomical template of experiment 1 were spatially transformed to match the group template space of experiment 2 using likewise the DARTEL approach. We report results at a level of p<0.001 uncorrected for multiple comparisons.

## 3 Results

### 3.1 Behavioral and Physiological Measures

#### 3.1.1 Experiment 1

Knee diameters were measured to determine the amount of joint swelling as an index of inflammation. Ipsilateral knee diameter in CFA pain model rats was significantly increased compared to vehicle treated rats at day 22 (mean KD CFA treated rats 10.8 ± 0.15 [SEM], mean KD control group 8.7mm ± 0.03, t(28)=14.2, p<0.001; Figure 1A). CFA pain model rats also showed decreased grip strength (mean GS CFA animals 931.0kg ± 30.1, mean GS control group 1259.0kg ± 10.6, t(28)=17.4, p<0.001; Figure 1B), weight on ipsilateral paw (mean DWB CFA treated rats 27.7% ± 1.5, mean DWB control group 49.5% ± 0.7, t(28)=13.9, p<0.001, Figure 1C) and amount of burrowed gravels (mean BB CFA treated rats 444.5g ± 121.5, mean BB control group 1238.2g ± 72.2, t(28)=5.8, p<0.001, Figure 1D). The results suggest that CFA pain model rats showed evoked and spontaneous inflammatory knee joint pain at the time of fMRI scanning.

**Figure 1:**
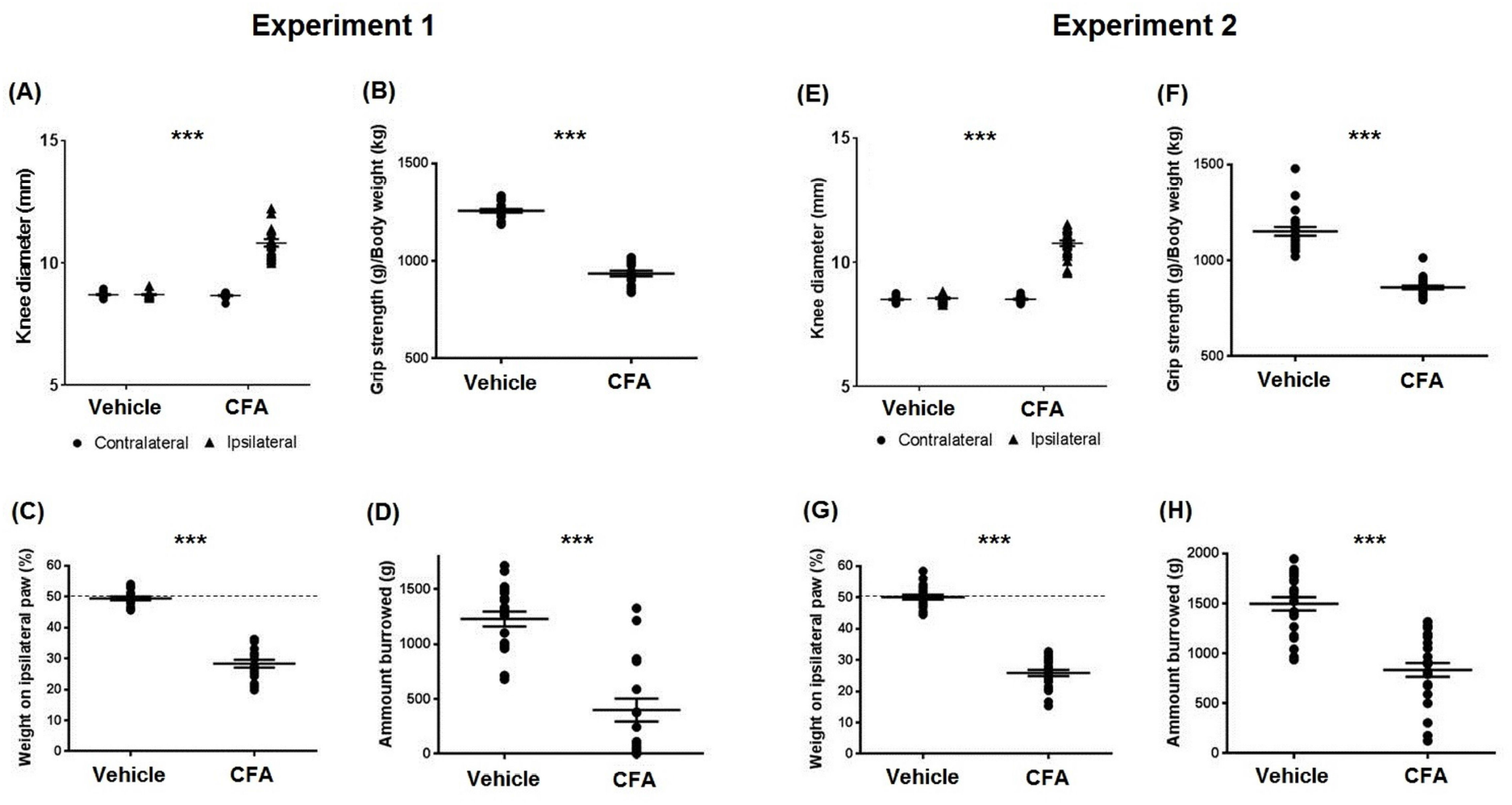
Behavioral and physiological changes of CFA pain model rats compared to controls at the time of fMRI scanning. (A-D) Experiment 1, (E-H) Experiment 2. (A)&(E) Changes in contralateral and ipsilateral knee diameter (KD) in vehicle and CFA rats at day 22 (in experiment 1) or the value of day 21 (in experiment 2). (B)&(F) Changes in hind limb grip strength (GS) in vehicle and CFA rats. Data is shown as an average value of day 18 and 22 (in experiment 1) or the value of day 20 (in experiment 2) expressed as grip strength/body weight. (C)&(G) Changes in dynamic weight bearing (WBD) on the ipsilateral paw in vehicle and CFA rats. Each data were shown as an average value of day 18 and 22 (in experiment 1) or the value of day 20 (in experiment 2). (D)&(H) Changes in burrowing behavior (BB) in vehicle treated and CFA pain model rats at day 18 (in experiment 1) or day 20 (in experiment 2). *** indicates p<0.001.

#### 3.1.2 Experiment 2

The replication experiment yielded very similar behavioral results. Ipsilateral knee diameter in CFA treated rats was significantly increased compared to sham injections at day 21 (mean KD CFA animals 10.7mm ± 0.11, mean KD control group 8.6mm ± 0.03, t(41)=18.6, p<0.001; Figure 1E). Likewise CFA pain model rats showed decreased grip strength (mean GS CFA animals 860.8kg ± 10.0, mean GS control group 1152.5kg ± 22.8, t(41)=11.9, p<0.001; Figure 1F), dynamic weight bearing on the ipsilateral paw (mean DWB CFA treated rats 25.8% ± 1.0, mean DWB control group 50.1% ± 0.8, t(41)=18.5, p<0.001; Figure 1G) and amount of burrowed gravels (mean BB CFA treated rats 868.5g ± 64.3, mean BB control group 1497.4g ± 67.0, t(41)=6.8, p<0.001; Figure 1H).

### 3.2 Resting State Functional Connectivity

#### 3.2.1 Experiment 1

The ACC seed region showed reduced functional connectivity in the CFA pain group compared to control animals with central parts of the contralateral somatosensory cortex and dorsal portions of the cingulate cortext (Fig. 2A). Additionally, the ACC exhibited increased functional connectivity in the pain group with rostral portions of the somatosensory cortex and in the subcortex with the bilateral striatum, structures of the basal forebrain region, hypothalamic region / preoptical area (POA) and the area of the bed nucleus of the stria terminalis (BNST; see Fig. 2A).

**Figure 2:**
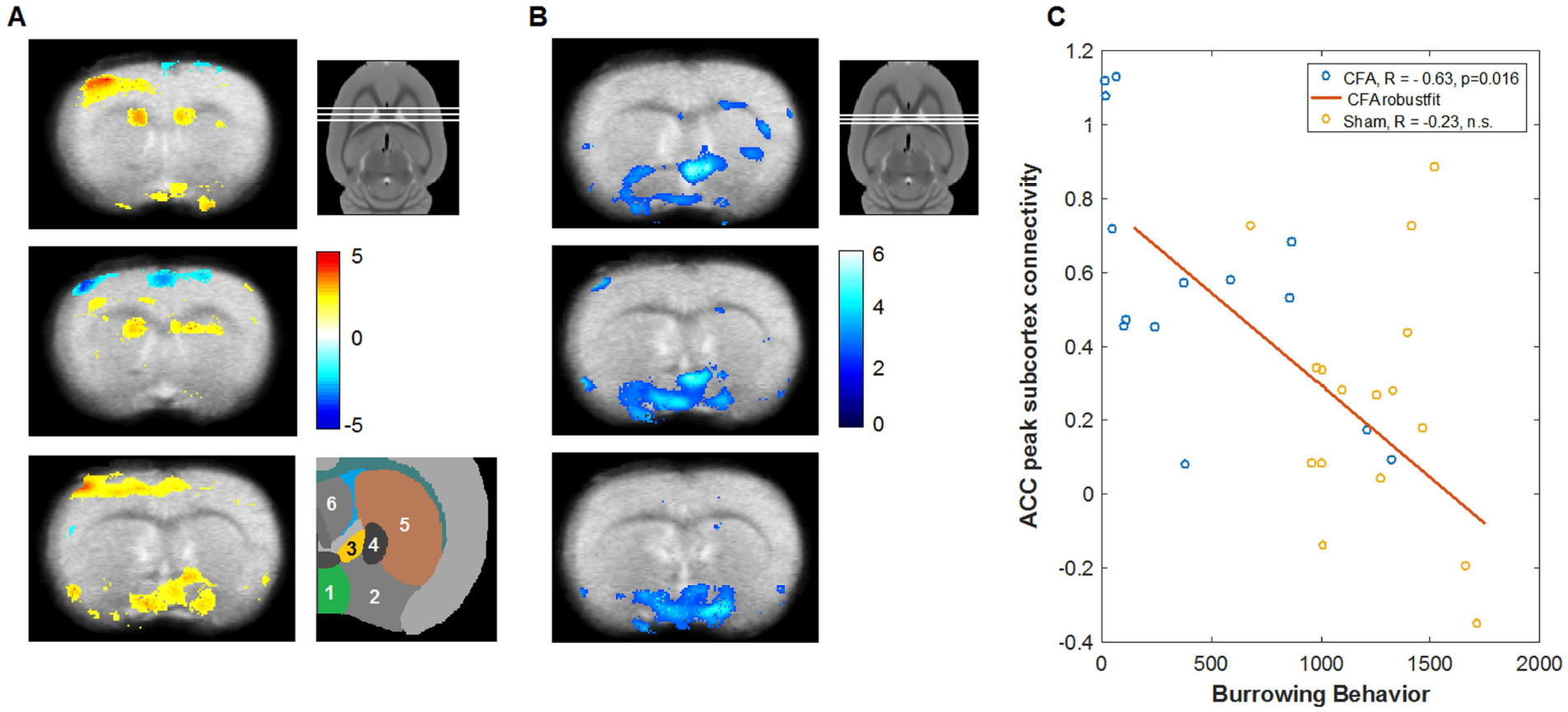
Changes in resting state functional connectivity in CFA pain model rats compared to control and correlation with burrowing behavior in experiment 1. (A) Changes in ACC resting state functional connectivity in CFA pain rats compared to vehicle treated animals. The ACC showed increased functional connectivity in CFA pain rats with parts of the contralateral somatosensory cortex (upper and lower panel), the bilateral striatum (upper and middle panel) and with the basal forebrain region / preoptical area and the region of the bed nucleus of the stria terminalis (lower panel). ACC functional connectivity was decreased in parts of the cingulate cortex itself and in the somatosensory cortex (middle panel). The 3 white lines in the smaller panel next to the upper panel of (A) indicate the approximate rostrocaudal position of the 3 axial sections of (A) in the rat brain. The color bar indicates t values. Hot colors represent increased functional connectivity and cold colors decreased connectivity. A schematic representation of the involved subcortical regions is shown next to the lower panel of (A). (1) Hypothalamic/preoptical area, (2) basal forebrain region, (3) bed nucleus of the stria terminalis, (4) globus pallidus, (5) striatum, (6) septal region. (B) Negative correlation of burrowing behavior with ACC functional connectivity. The color bar indicates t values. The visualization threshold is set to p<0.005 uncorrected in (A) and (B). Note the spacial overlap of increased ACC-subcortical connectivity in (A) with the negative connectivity-behavior correlation in (B) and the slightly different positions of the axial sections in (B) compared to (A). (C) Illustration of the negative correlation between burrowing behavior with the individual connectivity strength at the subcortical peak in the upper panel of (B). ACC connectivity shows a marked negative correlation with burrowing in the CFA pain group but not in the control group.

In CFA pain animals we found a marked negative correlation between ACC functional connectivity and innate burrowing behavior in the latter regions (POA & BNST; Fig. 2B), i.e. higher functional connectivity between ACC and these regions was associated with suppression of burrowing behavior. For illustration purposes, Figure 2C shows the correlation of connectivity strength between ACC with the peak subcortical region of interest and burrowing behavior. ACC connectivity shows a negative correlation with burrowing in the CFA pain group (r=-0.63, p=0.02) but not in the control group (r=-0.23, p=0.39). The interaction failed, however, significance (z=1.2, p=0.11).

We found no significant correlations between ACC connectivity with knee diameter, weight bearing and grip strength.

#### 3.2.2 Experiment 2

In experiment 2 we observed increased ACC functional connectivity in CFA pain animals compared to control in the somatosensory cortex, in the contralateral striatum in a region similar to experiment 1, and sporadically with a number of other cortical and subcortical regions (Fig. 3A). No regions with reduced ACC connectivity were observed in CFA pain rats compared to control. In accordance with experiment 1 we found a strong negative correlation between ACC-subcortical functional connectivity and the individual spontaneous burrowing behavior in the CFA pain group. In the basal forebrain region this correlation showed a close spatial relationship with the findings from the first experiment (Figure 3B) and can be likewise presumably attributed to the hypothalamic region and the adjacent BNST. Similar to the first experiment we exclusively found this negative correlation in this region in the CFA pain group (r=-0.74, p<0.001) but not in the healthy control group (r=-0.16, n.s., Figure 3C). The correlation between ACC connectivity and burrowing behavior was significantly more negative in CFA pain group compared to control animals (z=2.4, p=0.008). We also found a negative correlation of ACC functional connectivity and burrowing behavior in the thalamus and in the brainstem (Figure 3B). No significant correlations between ACC connectivity with knee diameter, weight bearing and grip strength and no positive correlation with burrowing behavior were observed.

**Figure 3:**
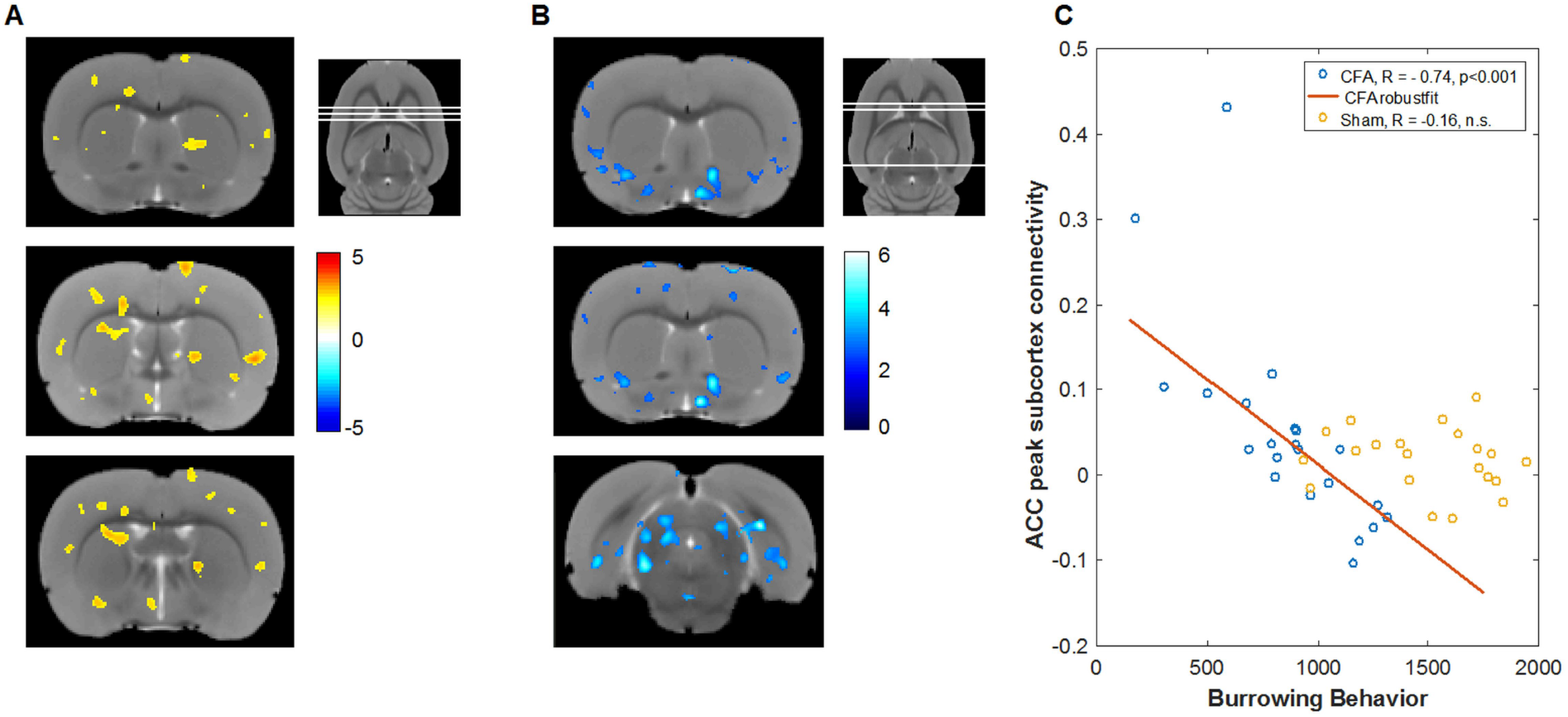
Changes in resting state functional connectivity in CFA pain model rats compared to control and correlation with burrowing behavior in experiment 2. (A) Changes in ACC resting state functional connectivity in CFA pain rats compared to vehicle treated animals. The ACC showed increased functional connectivity in CFA pain rats with the left dorsal striatum in a region similar to experiment 1 (middle and lower panel) and also with the right striatum (upper panel). Moreover, CFA animals exhibited increased ACC functional connectivity with a number of additional cortical and subcortical regions. No regions with reduced ACC connectivity were observed. To allow for comparability the colorbar range is kept identical to Figure 2. The visualization threshold is likewise set to p<0.005 uncorrected. The 3 white lines in the smaller panel next to the upper panel of (A) indicate the approximate rostrocaudal position of the 3 axial sections of (A) in the rat brain. (B) Negative correlation of burrowing behavior with ACC functional connectivity. On the individual level a negative correlation of burrowing with ACC connectivity strength was found similar to experiment 1 in the preoptical region and an area that can be most probably assigned to the bed nucleus of the stria terminalis (upper and middle panel). Moreover, we observed a negative correlation between burrowing and ACC connectivity strength in the brainstem (lower panel). (C) Negative correlation between burrowing behavior and the individual connectivity strength at the BNST peak (uppper and middle panel of B). ACC functional connectivity strength shows a strong negative correlation with burrowing behavior in the CFA animals in the BNST region that is significantly more negative compared to the control group.

## 4 Discussion

In two separate experiments we studied spontaneous burrowing behavior in the CFA model of persistent inflammatory arthritis pain and assessed ACC resting-state functional connectivity compared to controls. The behavioral measures consistently indicated inflammatory nociception related to the CFA-treated joint at the time of MRI measurement by showing ipsilateral swelling, reduced motoric ability and reduced weight bearing. As hypothesized we also observed reduced burrowing behavior accounting on average for less than 50% of the behavior in control animals. Resting-state fMRI in both experiments revealed a pronounced negative correlation between the individual strength of ACC functional connectivity and burrowing behavior in structures of the basal forebrain, comprising the hypothalamic region and the bed nucleus of the stria terminalis (BNST). No correlations with the ACC functional connectivity were observed for motor responses directly associated with joint pain. The findings therefore indicate a relatively specific relationship between ACC functional coupling and the suppression of burrowing behavior by persistent pain and suggest that ACC connectivity might be a good marker for the affective-motivational component of pain in rodents.

Across mammalian species neuronal responses to noxious stimuli have been shown to involve a number of brain regions including the primary and secondary somatosensory cortex, the ACC and the insular cortex (Iadarola et al., 1998; Lenz et al., 1998; Peyron, Laurent, and Garcia-Larrea, 2000; Ploner et al., 2002; Pohlmann et al., 2016; Thompson and Bushnell, 2012). Although the precise contribution of each region to the generation of pain remains to be elucidated, neuroimaging studies highlighted some general characteristics of the involved regions. While the somatosensory cortex responds faster to dimensional features of noxious stimuli, and is more engaged when larger body surfaces are exposed to pain, the ACC as part of the limbic system is thought to be modulated by the affective relevance pertinent to a change in motivational tone or response selection (Peyron, Laurent, and Garcia-Larrea, 2000; Ploner et al., 2002). The ACC is often conceptualized as a nexus for the processing of external salient stimuli, autonomic response regulation and subsequent affective learning (Gao et al., 2004; Vogt, 2005). For example, fear learning in mice through observing other mice receiving painful foot shocks has been demonstrated to involve the ACC (Jeon et al., 2010). In the context of pain, the ACC is thought to allow potentially harmful stimuli to engage appropriate affective and motivationally relevant behaviors, and as such one would expect the strength of ACC-based connectivity to be associated with elevated aversive behavior and reduced behaviors of well-being. Spontaneous burrowing behavior in rodents is considered as one of the latter (Deacon, 2006), and hence the association of higher connectivity between ACC and subcortex with reduced burrowing is consistent with a possible functional inhibitory pathway.

ACC connectivity showed increased functional coupling in the CFA group with a number of cortical and subcortical brain structures. Some of the regions such as the dorsal striatum and parts of the somatosensory cortex were found to consistently exhibit increased functional connectivity with the ACC across both experiments. However, a spatially more congruent picture between both data sets was observed regarding the association between the ACC-subcortex connectivity strength and the interindividual spectrum of spontaneous burrowing behavior. In both experiments we found in CFA-pain animal a similar negative correlation between burrowing and ACC connectivity in the basal forebrain region comprising hypothalamic / preoptical nuclei and the BNST, whereas no positive or negative correlation with burrowing was found in control animals in the brain. While the association between ACC-hypothalamic coupling and burrowing behavior might reflect a general homeostatic dimension related to the animal’s behavioral suppression, the correlation between burrowing and coupling strength with the BNST is especially interesting as it has been linked to sustained vigilance associated with ambiguous or distant threat cues. This potentially provides a direct link to the affective dimension of pain, and illustrates burrowing behavior as an expression of the affective-motivational tone of the animals. That is, when a limb is injured, the potential danger by predators and accordingly monitoring of potential threats becomes much more important, so that behaviors such as burrowing are no longer prioritized. In line with this lesions of the BNST have been demonstrated to disrupt the individual variability in the rodent’s anxiety-like behavior (Durvarci 2009). Finally, the role of the BNST as a site of integration of limbic forebrain information is also supported by tracing studies showing direct BNST connections with the ACC (area 32) (Kash et al., 2015).

The findings have implications for pain testing in animals. Most tests of pain in chronic pain models evaluate evoked pain, in which enhanced defensive behaviors are observed in response to stimulation of some sort. Such tests are likely to be highly sensitive to hyperalgesia and allodynia, but less so to the overall ‘wellbeing’ that accompanies it. Other tests, designed to capture persistent pain better, such as dynamic weight bearing, may still involve modulation of motor responses related to the pain associated with movement, but might be unlikely to relate purely to affective-motivational component of pain. In contrast, burrowing is an innate motivated behavior, and in the case of hind leg CFA injection, modulation by pain is more likely to reflect the underlying affective suppression of behavior in a way not directly related to exacerbation of the pain-inducing lesion. Given accumulating evidence that the transition to chronic pain can be characterized as the formation of a pathological affective state (Apkarian, Baliki, and Farmer, 2013) pain tests related to the affective-motivational dimension of the sustained pain state might be more closely related to relevant pathophysiological changes associated with disease progression. That suppression of burrowing behavior relates to ACC connectivity, which is strongly implicated in affective components of pain, further supports this notion, and adds to evidence that burrowing may provide a valuable complement to conventional measures in the evaluation of pain in rodent models of chronic pain. Assessing motivational parameters in the context of sustained pain in animals also captures information that has greater relevance to general well-being and higher-order biological goals compared to purely local measures, which is in line with the recent notion that more natural animal models might have a better translational validity (Klinck et al., 2017) and likewise with the finding that also in humans local parameters have only limited prognostic value.

Several limitations of the study should be noted. Experiments were conducted in anaesthetized animals and we cannot exclude that the narcotic agent interfered to a certain extent with the involved nociceptive mechanisms or the observed association between resting state connectivity and suppression of behavior. However, even high concentrations of isoflurane exert only mild antinociceptive effects and reexamination of the behavioral measures after fMRI measurements showed no significant changes. Anesthetics naturally interfere with the brain-wide spatiotemporal organization of functional networks, but it appears very unlikely that a narcotic agent selectively induced associations between spontaneous fluctuations of resting-state brain activity and behavioral measures in one group of animals.

Notwithstanding, neuroimaging experiments alone are inherently correlational, and it cannot be assumed that the ACC connections mediate a direct causal suppression of affective-motivational behavior on the basis of the evidence provided. Even if the connectivity ***is*** causal, one needs to remain cautious about generalizing to other motivated behaviors, in case the pathway was specific to burrowing behavior. With this in mind, it is difficult to find an exact parallel to burrowing behavior in humans. However, these issues can be potentially addressed. For instance, the suppression of affective and motivated behaviors could be studied in human pain patients with regard to ACC connectivity. The employed experimental paradigm also provides interesting perspectives for future studies in rodents. Pharmacological studies using systemic analgesics could investigate correlations between suppression / reinstatement of burrowing behavior, ACC connectivity and the development of the painful condition. And targeted manipulations of the ACC (local pharmacology, optogenetics) with concurrent neuroimaging could be highly informative with regard to the identification of potential causal relationships.

## Acknowledgements

Masahide Fujita gave advice for animal transportation.

## References

Andrews, N. et al. (2012). “Spontaneous burrowing behaviour in the rat is reduced by peripheral nerve injury or inflammation associated pain”. In: Eur J Pain 16.4, pp. 485–95.

Apkarian, A. V., M. N. Baliki, and M. A. Farmer (2013). “Predicting transition to chronic pain”. In: Current opinion in neurology 26.4, pp. 360–367.

Ashburner, J. (2007). “A fast diffeomorphic image registration algorithm”. In: Neuroimage 38.1, pp. 95–113.

Büchel, C., K. Bornhövd, M. Quante, V. Glauche, B. Bromm, and C. Weiller (2002). “Dissociable Neural Responses Related to Pain Intensity, Stimulus Intensity, and Stimulus Awareness within the Anterior Cingulate Cortex: A Parametric Single-Trial Laser Functional Magnetic Resonance Imaging Study”. In: Journal of Neuroscience 22.3, pp. 970–976.

Calejesan, A. A., S. J. Kim, and M. Zhuo (2000a). “Descending facilitatory modulation of a behavioral nociceptive response by stimulation in the adult rat anterior cingulate cortex”. In: European Journal of Pain 4.1, pp. 83–96.

Calejesan, A. A., S. J. Kim, and M. Zhuo (2000b). “Descending facilitatory modulation of a behavioral nociceptive response by stimulation in the adult rat anterior cingulate cortex”. In: European Journal of Pain 4.1, pp. 83–96.

Deacon, R. (2006). “Burrowing in rodents: a sensitive method for detecting behavioral dysfunction”. In: NATURE PROTOCOLS-ELECTRONIC EDITION- 1.1, p. 118.

Eto, K., H. Wake, M. Watanabe, H. Ishibashi, M. Noda, Y. Yanagawa, and J. Nabekura (2011). “Inter-regional contribution of enhanced activity of the primary somatosensory cortex to the anterior cingulate cortex accelerates chronic pain behavior”. In: Journal of Neuroscience 31.21, pp.7631–7636.

Friston, K. J. (1994). “Functional and effective connectivity in neuroimaging: A synthesis”. en. In: Human Brain Mapping 2.1-2, pp. 56–78.

Gao, Y.-J., W.-H. Ren, Y.-Q. Zhang, and Z.-Q. Zhao (2004). “Contributions of the anterior cingulate cortex and amygdala to pain-and fear-conditioned place avoidance in rats”. In: Pain 110.1, pp. 343–353.

Iadarola, M. J., K. F. Berman, T. A. Zeffiro, M. G. Byas-Smith, R. H. Gracely, M. B. Max, and G. J. Bennett (1998). “Neural activation during acute capsaicin-evoked pain and allodynia assessed with PET.” In: Brain 121.5, pp. 931–947.

Jeon, D., S. Kim, M. Chetana, D. Jo, H. E. Ruley, S.-Y. Lin, D. Rabah, J.-P. Kinet, and H.-S. Shin (2010). “Observational fear learning involves affective pain system and Cav1.2 Ca2+ channels in ACC | Nature Neuroscience”. eng. In: Nature Neuroscience 13.4, pp. 482–488.

Kash, T. L., K. E. Pleil, C. A. Marcinkiewcz, E. G. Lowery-Gionta, N. Crowley, C. Mazzone, J. Sugam, J. A. Hardaway, and Z. A. McElligott (2015). “Neuropeptide Regulation of Signaling and Behavior in the BNST”. In: Molecules and Cells 38.1, pp. 1–13.

Klinck, M. P., J. S. Mogil, M. Moreau, B. D. X. Lascelles, P. A. Flecknell, T. Poitte, and E. Troncy (2017). “Translational pain assessment: could natural animal models be the missing link?” In: PAIN 158.9, p. 1633.

Kundu, P., M. D. Santin, P. A. Bandettini, E. T. Bullmore, and A. Petiet (2014). “Differentiating BOLD and non-BOLD signals in fMRI time series from anesthetized rats using multi-echo EPI at 11.7T”. In: NeuroImage 102, pp. 861–874.

Lenz, F., M. Rios, A. Zirh, D. Chau, G. Krauss, and R. Lesser (1998). “Painful stimuli evoke potentials recorded over the human anterior cingulate gyrus”. In: Journal of neurophysiology 79.4, pp. 2231–2234.

Lloyd, D., G. Di Pellegrino, and N. Roberts (2004). “Vicarious responses to pain in anterior cingulate cortex: is empathy a multisensory issue?” In: Cognitive, Affective, & Behavioral Neuroscience 4.2, pp. 270–278.

Moisset, X. and D. Bouhassira (2007). “Brain imaging of neuropathic pain”. In: Neuroimage 37, S80–S88.

Peyron, R., B. Laurent, and L. Garcia-Larrea (2000). “Functional imaging of brain responses to pain. A review and meta-analysis (2000)”. In: Neurophysiologie Clinique/Clinical Neurophysiology 30.5, pp. 263–288.

Ploner, M., J. Gross, L. Timmermann, and A. Schnitzler (2002). “Cortical representation of first and second pain sensation in humans”. In: Proceedings of the National Academy of Sciences 99.19, pp.12444–12448.

Pohlmann, A. et al. (2016). “Normothermic Mouse Functional MRI of Acute Focal Thermostimulation for Probing Nociception”. En. In: Scientific Reports 6, p. 17230.

Price, D. D. (2000). “Psychological and neural mechanisms of the affective dimension of pain”. In: Science 288.5472, pp. 1769–1772.

Rainville, P. (2002). “Brain mechanisms of pain affect and pain modulation”. In: Current Opinion in Neurobiology 12.2, pp. 195–204.

Rainville, P., G. H. Duncan, D. D. Price, B. Carrier, and M. C. Bushnell (1997). “Pain affect encoded in human anterior cingulate but not somatosensory cortex”. In: Science 277.5328, pp. 968–971.

Reichman, O. and S. C. Smith (1990). “Burrows and burrowing behavior by mammals”. In: Current mammalogy 2, pp. 197–244.

Seminowicz, D. A., A. L. Laferriere, M. Millecamps, S. Jon, T. J. Coderre, and M. C. Bushnell (2009). “MRI structural brain changes associated with sensory and emotional function in a rat model of long-term neuropathic pain”. In: Neuroimage 47.3, pp. 1007–1014.

Singer, T., B. Seymour, J. O’doherty, H. Kaube, R. J. Dolan, and C. D. Frith (2004). “Empathy for pain involves the affective but not sensory components of pain”. In: Science 303.5661, pp.1157–1162.

Thompson, S. J. and M. C. Bushnell (2012). “Rodent functional and anatomical imaging of pain”. In: Neuroscience Letters. Neuroimaging of Pain: Insights into normal and pathological pain mechanisms 520.2, pp. 131–139.

Valdés-Hernández, P. A., A. Sumiyoshi, H. Nonaka, R. Haga, E. Aubert-Vásquez, T. Ogawa, Y. Iturria-Medina, J. J. Riera, and R. Kawashima (2011). “An in vivo MRI Template Set for Morphometry, Tissue Segmentation, and fMRI Localization in Rats”. In: Frontiers in Neuroinformatics 5.

Vogt, B. A. (2005). “Pain and emotion interactions in subregions of the cingulate gyrus”. In: Nature Reviews Neuroscience 6.7, pp. 533–544.

Vogt, B. A., D. M. Finch, and C. R. Olson (1992). “Functional heterogeneity in cingulate cortex: the anterior executive and posterior evaluative regions”. In: Cerebral Cortex, pp. 435–443.

Vogt, B. A. and A. Peters (1981). “Form and distribution of neurons in rat cingulate cortex: Areas 32, 24, and 29”. In: The Journal of Comparative Neurology 195.4, pp. 603–625.

Zhuo, M. (2007). “Neuronal mechanism for neuropathic pain”. In: Molecular pain 3.1, p. 14.

